# The Synergy between a Silver-Ruthenium Antimicrobial and aminoglycosides is based on severe macromolecular damage

**DOI:** 10.64898/2025.12.21.695785

**Authors:** Emmanuel P. Oladokun, Gracious Y. Donkor, Julius K. Narh, Grady D. Jacobson, Cade Ward, Patrick O. Tawiah, Lisa K. Polzer, Kevin A. Edwards, Kyle A. Floyd, Jan-Ulrik Dahl

**Affiliations:** School of Biological Sciences, Illinois State University, Normal, IL

## Abstract

The rise of multidrug-resistant (MDR) bacterial pathogens, including uropathogenic *Escherichia coli* (UPEC), highlights the urgent need for alternative treatment strategies to restore antibiotic efficacy. The silver-ruthenium antimicrobial AGXX^®^ exerts potent bactericidal effects through the production of reactive oxygen species (ROS); however, its potential synergy with antibiotics has not been thoroughly investigated. Here, we show that sublethal concentrations of AGXX^®^ strongly enhance aminoglycoside-mediated killing across a diverse panel of Gram-negative and Gram-positive MDR clinical isolates, including highly aminoglycoside-resistant strains. Combinational treatments significantly reduced the effective concentrations of gentamicin, tobramycin, kanamycin, and amikacin required for bacterial killing. Mechanistic analyses revealed that AGXX^®^/aminoglycoside co-treatment induces pronounced intracellular ROS accumulation, resulting in severe proteotoxic stress, extensive protein aggregation, and DNA damage. Scavenging ROS abolished synergistic killing, establishing oxidative imbalance as the primary driver of the synergy between both antimicrobials. We further identify polyphosphate as a key bacterial defense mechanism that mitigates ROS accumulation, proteotoxicity, and genotoxic stress during combinational treatment. Moreover, AGXX^®^–aminoglycoside synergy was preserved in an artificial urine medium and across clinical UPEC isolates, underscoring its relevance to urinary tract infections. Together, these findings position AGXX^®^ as a potent aminoglycoside adjuvant that restores antibiotic efficacy through ROS-driven macromolecular damage, supporting its development for combination therapies against MDR bacterial infections.

## INTRODUCTION

Originally isolated from *Streptomyces spp.* and *Micromonaspora*, respectively, aminoglycoside antibiotics have now also been developed as semisynthetic derivatives (1). These bactericidal antimicrobials are polycationic and enter bacteria in three steps. Firstly, establishing ionic interactions between the polycationic moieties of aminoglycosides and negatively charged components of the cell surface, including those found in lipopolysaccharide (LPS) (2, 3) displaces divalent cations from aminoglycoside binding sites. This, in turn, results in the destabilization of LPS components and the disruption of outer membrane integrity in an energy-independent way (4). Driven by an increased proton motive force, particularly in aerobically respiring bacteria, aminoglycosides cross the plasma membrane and enter the cytoplasm, where they bind to the aminoacyl-tRNA recognition site and interact with the 16S ribosomal RNA, ultimately causing protein mistranslation that can lead to protein misfolding [recently reviewed in (1)]. Misfolded proteins can be inserted into the plasma membrane and form irregular membrane channels that further exacerbate aminoglycoside uptake and subsequent bactericidal effects (5, 6). Although controversial (7), ROS production resulting from hyperactivation of the electron transport chain in aerobically respiring bacteria has been proposed as a potential contributor to the bactericidal activity of aminoglycosides (5, 6, 8). Likewise, aminoglycoside resistance observed in most anaerobic bacteria could be explained by a lack of sufficiently high proton motive force that aminoglycoside uptake depends on (5, 9). Moreover, Ezraty *et al.* showed that iron-sulfur cluster-containing proteins play an important role in aminoglycoside uptake, ultimately contributing to the bactericidal effects of these antibiotics (10).

Resistance to aminoglycosides has now been reported in all ESKAPEE pathogens, including the Gram-negative bacterium *Escherichia coli*. *E. coli* colonizes the gastrointestinal (GI) tract of human newborns shortly after birth, where they establish themselves as part of the commensal gut microbiome. Beyond commensal symbiosis, additional *E. coli* serotypes can be pathogenic, resulting in life-threatening diarrheal disease, including enteropathogenic, enterohemorrhagic, enterotoxigenic, enteroinvasive, enteroaggregative, and diffusely adherent *E. coli* serotypes (11). Moreover, extraintestinal pathogenic *E. coli* serotypes (ExPEC) can often still colonize the GI tract and serve commensal functions. It is only after dissemination from the GI tract and entry into their primary site of infection that ExPEC strains become pathogenic. The most common ExPEC serotype is uropathogenic *E. coli* (UPEC), the etiologic agent of most urinary tract infections (UTIs) (11). UPEC traverse the urethra to establish infection within the bladder (cystitis) and can further ascend the ureters to cause infection within the kidneys (pyelonephritis), which, if left untreated, can result in the development of potentially life-threatening septicemia (12, 13). UTIs account for approximately 20% – 50% of all nosocomial infections (14), many of which occur in catheterized patients. Biofilm formation on in-dwelling urinary catheters is the main driving force behind catheter-associated UTIs (CAUTIs) and serves as a reservoir for continued seeding of infection within the urinary tract (12, 13, 15). There is also increasing evidence of the presence of antimicrobial resistance (AMR)- related genes in UPEC (16). The inherent infection mechanisms of UPEC, the ability to form catheter-associated biofilms, and the rise in prevalence of AMR highlight the urgent need for the development of improved antimicrobial compounds and catheter surface coatings to treat or prevent UTI (17, 18).

Despite its long-standing history and high efficacy against bacteria, the antimicrobial mode of action of silver (Ag) remains poorly understood, although changes in DNA condensation, membrane alteration, and protein damage have been associated with Ag (19). Ag^+^ ions interact with cysteine thiols, destabilize iron-sulfur clusters, replace metal-containing cofactors, and and elicit reactive oxygen species (ROS) generation, thereby damaging a wide range of proteins (19). Over the last two decades, silver derivatives have received increased attention for use as antimicrobial surface coatings that can protect from biofilm-forming bacteria and reduce the risk of nosocomial infections (20). One such silver-containing surface coating is AGXX^®^ (21, 22). All AGXX^®^ formulations consist of a silver-ruthenium complex, but differ in the silver-ruthenium ratio, particle size, and production procedure, which may affect their antimicrobial activity (23, 24). AGXX^®^ has been shown to be more potent than silver alone, inhibiting the growth of a range of pathogenic bacteria by eliciting a thiol-specific oxidative stress response (22, 25–27). We recently reported that AGXX^®^ potentiates the efficacy of aminoglycoside antibiotics, when used in combination, against the opportunistic pathogen *Pseudomonas aeruginosa* (26). The synergistic bactericidal effects against *P. aeruginosa* are mediated by elevated ROS levels in the cells, which disrupt Fe/S-clusters in metabolic enzymes, lead to elevated free iron, trigger hydroxyl radical formation, and result in a significant increase in outer and inner membrane permeability to exacerbate aminoglycoside influx into the cell (26).

In this study, we provide novel insights into the cellular consequences of the synergy between AGXX^®^ and aminoglycosides across a variety of Gram-positive and Gram-negative pathogens, including UPEC and highly aminoglycoside-resistant clinical isolates of various ESKAPEE pathogens. This synergy appears to primarily rely on the generation of ROS, as co-supplementation with an antioxidant abolishes AGXX^®^/aminoglycoside-induced cell death. For the first time, we provide direct evidence that AGXX^®^/aminoglycoside co-treatments promote extensive protein aggregation and induce DNA damage. Despite the potent toxicity of AGXX^®^/aminoglycoside co-treatment, we found that UPEC can activate sophisticated defenses by expressing molecular chaperones, such as IbpA, and by producing the chemical chaperone polyphosphate (polyP). We observed that polyP reduces intracellular ROS accumulation, thereby decreasing DNA damage and protein aggregation, and positively affects UPEC survival, underscoring the significant role of polyP in oxidative stress defense.

## RESULTS

### AGXX^®^ enhances the bactericidal effects of aminoglycoside antibiotics in various multidrug-resistant pathogens

The rapid emergence of multidrug-resistant pathogens (MDR) requires alternative treatment options, including therapies that enhance the effectiveness of existing antibiotics. The silver-ruthenium antimicrobial AGXX^®^, which was shown to be effective against laboratory strains of both Gram-positive and Gram-negative pathogens (22–25, 27, 28), may provide such an opportunity. As previously reported by us, AGXX^®^ potentiates the activity of aminoglycoside antibiotics in the laboratory *P. aeruginosa* strain PA14, which is due to elevated aminoglycoside uptake as a result of ROS-induced membrane damage (26). To exclude strain-specific effects and assess the effectiveness of AGXX^®^/aminoglycoside combinations against other bacterial pathogens, we examined the impact of AGXX^®^/aminoglycosides against a panel of Gram-negative and Gram-positive clinical isolates obtained from burn-wound patients. With minimal inhibitory concentrations (MIC) between 1 and >2,000 μg/mL, these isolates differ quite significantly in their susceptibility to gentamycin (Gm) and tobramycin (Tob), respectively (**Supplementary Table S1**). We exposed *P. aeruginosa* strains 1, 36, and 40; *Klebsiella pneumoniae* strain 18; *Acinetobacter baumannii* strain 99; and *Staphylococcus aureus* strain 3 to the indicated sublethal concentrations of AGXX^®^720C, aminoglycosides, or their combination and monitored bacterial survival for 4 hours by enumerating colony-forming units (CFUs). Individual treatments with sublethal concentrations of AGXX^®^720C or aminoglycoside had minimal (*P. aeruginosa* strains 36 and 40) to no bactericidal effects across these strains (**Fig. 1A–G**). In contrast, combinational treatments with AGXX^®^720C and aminoglycosides resulted in rapid synergistic killing, with ∼3-log or greater reductions in CFUs observed in all tested strains after 4 hours. This effect was particularly pronounced against *P. aeruginosa* strain 1 and *A. baumannii* strain 99, which both have Gm/Tob MICs >500 μg/mL (**Supplementary Table S1**). Notably, sub-MIC concentrations of Gm/Tob, combined with AGXX^®^720C, significantly reduced survival by more than 3-log, thereby rendering these highly resistant isolates sensitive to aminoglycosides again (**Fig. 1C, 1E**). These findings demonstrate the therapeutic promise of AGXX^®^ as a broad-spectrum compound capable of enhancing aminoglycoside activity against both Gram-negative and Gram-positive MDR pathogens.

**Fig. 1.**
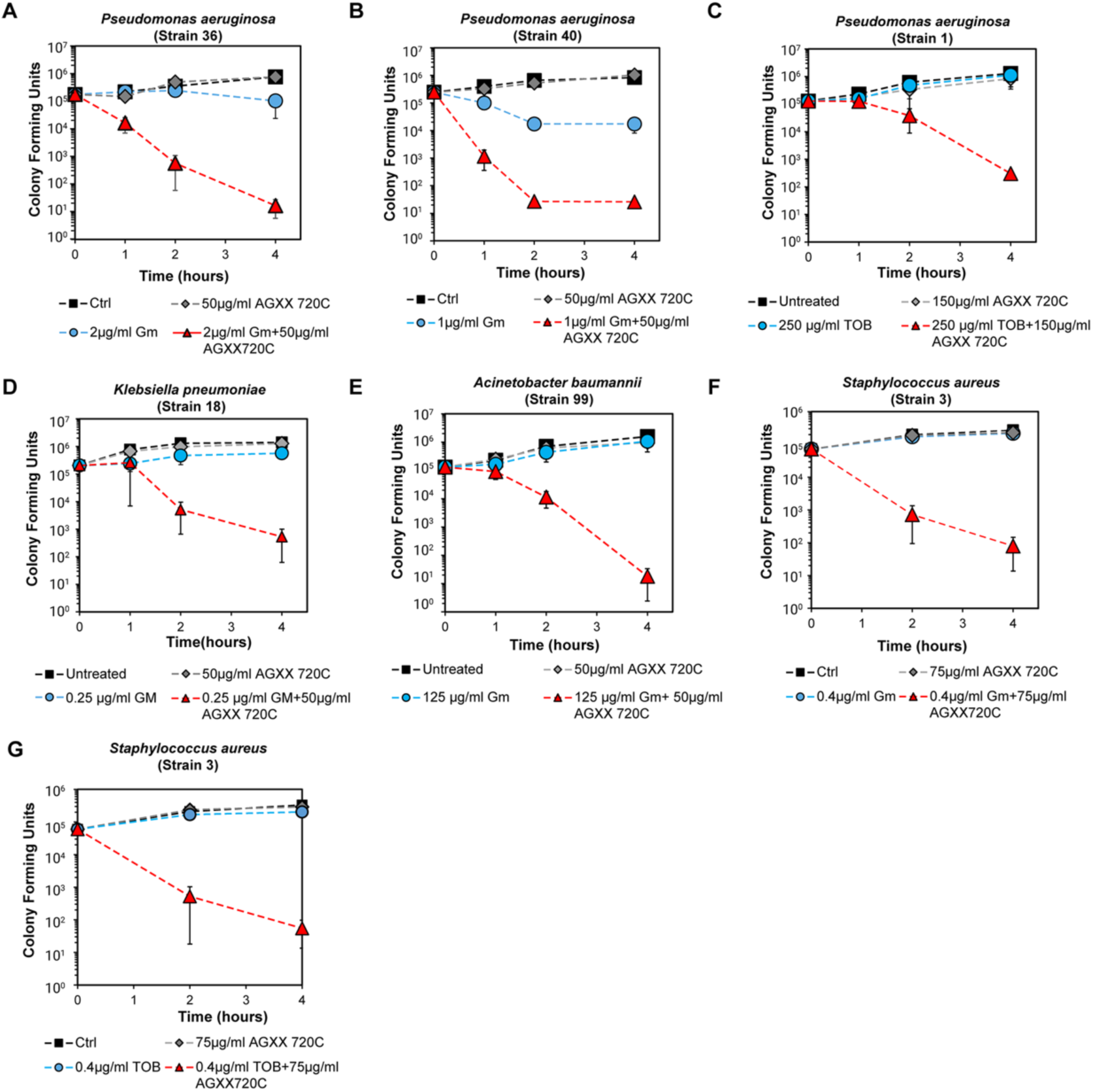
AGXX^®^ enhances the bactericidal effects of aminoglycoside antibiotics in various multidrug-resistant ESKAPEE pathogens. Time-killing survival studies were performed on various clinical isolates of *P. aeruginosa* (**A–C**), *K. pneumoniae* (**D**), *A. baumannii* (**E**), and *S. aureus* (**F, G**). Mid-log phase cells were exposed to sublethal concentrations of gentamycin (Gm), tobramycin (TOB), AGXX^®^720C, or combinations as indicated. Bacterial survival was quantified by determining colony-forming units (CFUs) at the indicated time points post-treatment (0–4 h). (n = 3, ±SD).

### AGXX^®^/aminoglycoside co-treatment causes severe oxidative stress in uropathogenic *E. coli* (UPEC)

Silver and silver-containing compounds have become increasingly popular as surface coatings on medical devices, such as catheters, to prevent bacterial attachment and reduce the risk of developing CAUTIs, a significant health problem for hospitalized patients and those living in long-term care facilities (15). The most prominent causative agent of CAUTIs is UPEC, which readily adheres to catheters and forms stress-/treatment-resistant biofilms. To examine the efficacy of AGXX^®^/aminoglycoside combinational treatment of UPEC, we treated mid-log phase CFT073 cells with sublethal concentrations of Tob, amikacin (Ami), Gm, or kanamycin (Kan), alone and in combination with 30 µg/mL of a more recent AGXX^®^ formulation (*i.e.,* AGXX^®^394C), and quantified bacterial survival via CFU enumeration after 180 minutes. Individual treatments of AGXX^®^394C and the four aminoglycosides had minimal effects on CFT073, only reducing survival by less than 1-log (**Fig. 2A**). However, combination treatment of AGXX^®^394C and any aminoglycoside (most notably Tob and Ami), decreased CFT073 survival by up to 4-logs (**Fig. 2A**). A similar synergistic effect was observed for AGXX^®^394C/Tob treatment of two UPEC clinical isolates VUTI156 (pyelonephritis) and VUTI207 (cystitis) after 60 minutes, which further demonstrates combinatorial effects with clinically relevant UPEC isolates (**Fig. 2B**). These experiments were conducted in artificial urine media (AUM), to more closely mimic conditions found within the urinary tract (29). Next, we sought to characterize the strength of the synergistic interaction between AGXX^®^ and aminoglycosides in CFT073. We performed a checkerboard assay at varying concentrations of Gm, Kan, Ami, and Tob, with or without 30 µg/mL AGXX^®^720C. We observed that the addition of AGXX^®^720C reduced the aminoglycoside concentrations required for complete growth inhibition by 16.7-fold for Gm, 3-fold for Kan, 4-fold for Ami, and 3.8-fold for Tob (**Supplementary Figure S1**), indicating that AGXX^®^ potentiates aminoglycoside antibiotics, independent of the type of aminoglycoside and/or AGXX^®^ formulation used.

**Fig. 2.**
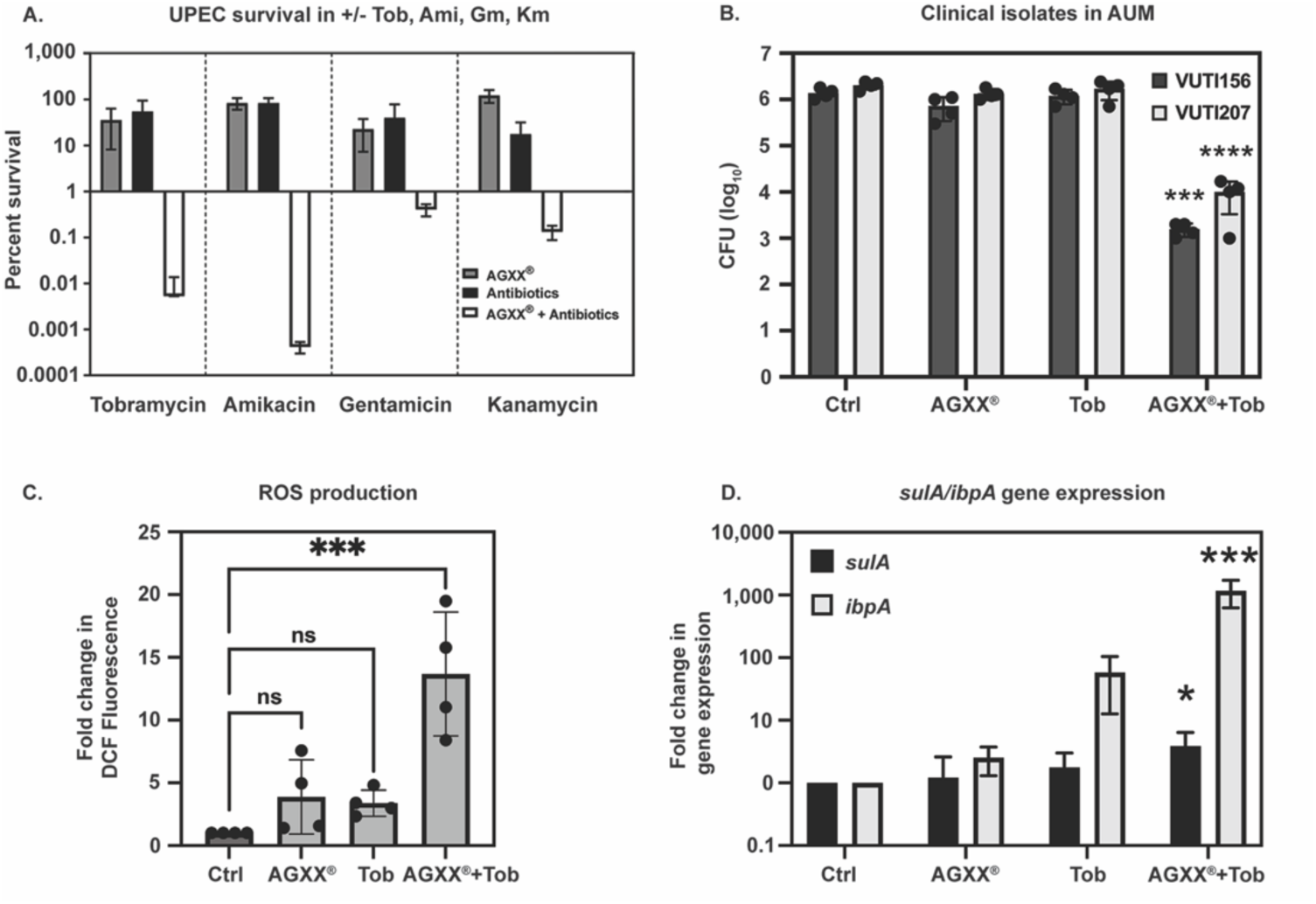
AGXX®/aminoglycoside co-treatment causes severe oxidative stress and elicits stress responses in UPEC. **(A)** Exponentially growing cells of the UPEC strain CFT073 were exposed to sublethal concentrations of tobramycin (Tob, 0.75 µg/mL), amikacin (Ami, 0.875 µg/mL), gentamicin (Gm, 0.01 µg/mL), kanamycin (Kan, 1 µg/mL), or AGXX^®^394C (30 µg/mL), individually and in combination. CFU counts were determined after 3 hours of incubation. The % survival, normalized to untreated CFUs, is plotted (n = 3-4, ±SD). **(B)** UPEC clinical isolates VUTI207 and VUTI156 were grown in artificial urine media (AUM) and treated with 1 µg/mL Tob or 30 µg/mL AGXX^®^394C individually and in combination. CFU counts were determined after 1 hour of incubation (n = 4, ±SD; statistical analysis: One-way ANOVA, Šídák’s multiple comparison test [compared to untreated controls], *** P < 0.001, **** P < 0.0001). (**C-D**) Mid-log phase CFT073 cells were treated with sublethal concentrations of Tob (0.5 µg/mL) or AGXX^®^394C (30 µg/mL) individually and in combination. **(C)** ROS levels were measured by H2DCFDA fluorescence (n = 4, ±SD, statistical analysis: One-way ANOVA, Dunnett’s posttest, ns = P > 0.05, *** P < 0.001). (**D**) *ibpA* and *sulA* transcript levels were determined by qRT-PCR. Expression of each gene was normalized to the housekeeping gene *rrsD* and calculated as fold change relative to untreated cells (n = 4, ±SD; statistical analysis: One-way ANOVA, Šídák’s multiple comparison test [compared to untreated controls], * P < 0.05, *** P < 0.001).

In *P. aeruginosa*, we previously showed that the synergistic effect between AGXX^®^ and sub-lethal aminoglycoside concentrations is attributed to increased accumulation of ROS (26). We observed a similar ROS accumulation in UPEC strain CFT073, where neither AGXX^®^394C nor Tob alone increased ROS levels, but AGXX^®^394C co-treatment with a sub-lethal Tob concentration resulted in a ∼13-fold increase in ROS levels relative to untreated controls (**Fig. 2C**). Increased ROS levels pose a risk to cellular macromolecules, especially proteins and nucleic acids (30, 31). However, bacteria have evolved intricate systems to deal with the negative consequences of ROS and eliminate ROS-mediated damage, including through transcriptional upregulation of heat-shock and SOS responses (32–36). To determine if AGXX^®^394C/Tob co-treatment elicits increased responses indicative of macromolecular damage, we examined the transcript levels of hallmark genes for protein aggregation (*i.e., ibpA*) and DNA damage (*i.e., sulA*) using qRT-PCR. Expression of *ibpA*, encoding a small heat shock protein that protects against proteotoxic stress (37), was induced 2.5-fold and 58-fold by individual exposure to AGXX^®^394C or Tob, respectively, while no significant change in expression of the gene *sulA*, a gene encoding for the SOS cell division inhibitor activated during DNA damage (38), was observed with individual treatments (**Fig. 2D**). However, AGXX^®^394C/Tob co-treatment resulted in over 1,000-fold increase in *ibpA* and a 5-fold increase in *sulA* expression (**Fig. 2D**), suggesting that the synergy between AGXX^®^ and aminoglycosides in CFT073 may be based on a more pronounced macromolecular damage.

### The synergy between AGXX^®^ and aminoglycosides is due to increased proteotoxic effects, evoking elevated expression of molecular chaperones

Following up on our transcriptional data pointing towards increased macromolecular damage in bacteria treated with AGXX^®^394C/Tob combinations, we assessed the extent of protein aggregate formation *in vivo* in the presence and absence of individual or combined treatments with sublethal AGXX^®^394C and/or Tob concentrations. We utilized a reporter strain harboring a chromosomal fusion between *ibpA* and monomeric superfolder green fluorescent protein under the control of the native *ibpA* promoter. IbpA expression is induced under stress conditions that induce protein unfolding (39), and IbpA-msfGFP has been used previously to monitor and quantify protein aggregation *in vivo* (27, 40). We exposed *E. coli* cells expressing IbpA-msfGFP to sublethal concentrations of AGXX^®^394C (15 µg/mL), Tob (0.3 µg/mL), or their combination for 1 hour, and then measured IbpA-msfGFP fluorescence by flow cytometry. As a positive control, we induced protein aggregation via heat shock by exposing cells to 50°C for 60 min, which resulted in the most pronounced IbpA-msfGFP fluorescence (**Supplementary Fig. S2**). The combination treatment of AGXX^®^394C and Tob induced a substantial shift in fluorescence intensity representing a ∼2-fold increase in IbpA-msfGFP fluorescence **(Fig. 3A-B)**, comparable to the 50°C heat-shock control, while there was no significant change in IbpA-msfGFP fluorescence observed for the individual treatments and untreated controls **(Fig. 3A-B)**. Visualization of IbpA-msfGFP fluorescent foci by confocal microscopy showed a similar trend for cells exposed to either individual or combined treatments for 60 minutes (**Fig. 3C-D**). Approximately 25% and 30% of cells showed a single IbpA-msfGFP focus per cell upon individual treatment with AGXX^®^394C or Tob, respectively, which is comparable to the 20% of cells with a foci observed in untreated cells (**Fig. 3C-D**). However, IbpA-msfGFP foci formation was significantly higher upon AGXX^®^394C/Tob combinational treatment, where 85% of cells had foci, and approximately 30% of those cells harbored multiple foci per cell (**Fig. 3C-D**). Overall, our results provide direct evidence for increased proteotoxicity as a result of the synergy between AGXX^®^394C and Tob.

**Fig. 3.**
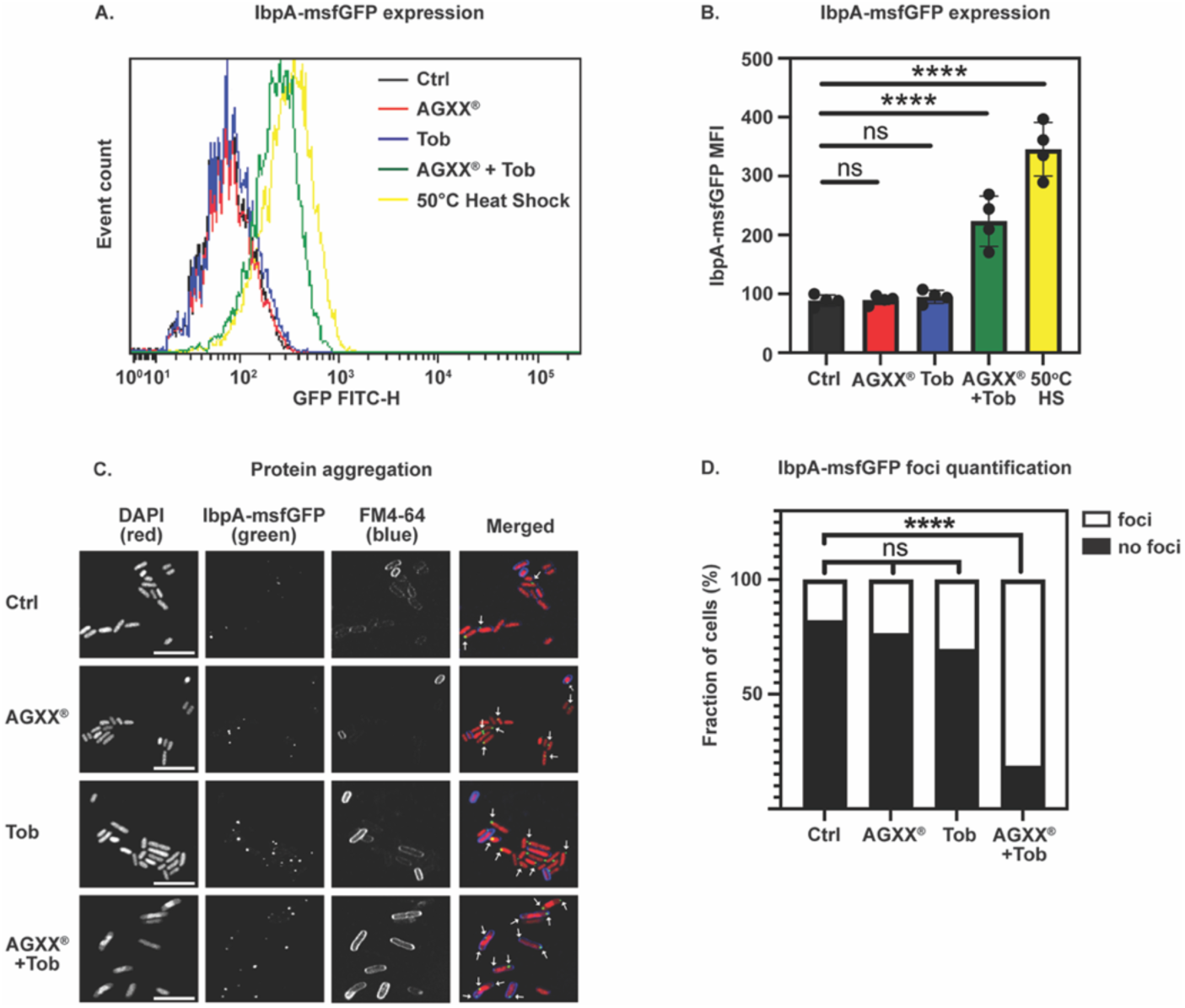
The synergy between AGXX^®^ and aminoglycosides is due to increased proteotoxic effects, evoking elevated expression of molecular chaperones. Cellular IbpA-msfGFP fluorescence was quantified after 1 hour of exposure of mid-log phase *E. coli* cells to sublethal concentrations of AGXX^®^394C (15 µg/mL), Tob (0.3 µg/mL), or their combination. Untreated cells serve as the negative control, and 50°C heat-shocked cells as the positive control. (**A**) Total cellular IbpA-msfGFP fluorescence determined by flow cytometry. One representative dataset of four biological replicates is shown (scale bar: 7.5 µm). (**B**) Quantification of mean fluorescence intensity (MFI) of IbpA-msfGFP fluorescence determined by flow cytometry (n = 4, ±SD; statistics: One-way ANOVA, Dunnett’s multiple comparison test, ns = P > 0.05, ****P < 0.0001). (**C**) For visualization and quantification of protein aggregates *in vivo*, cells were stained with DAPI (to visualize nucleic acids, red) and FM4-64 (to visualize membranes, blue). White arrows indicate foci formed by IbpA-msfGFP (green) binding to protein aggregates *in vivo*. Images representative of three biological replicates. (**D**) Quantification of the number of foci per cell from microscopy images as shown in **(C)** (n=3, statistics: Two-way Annova, Tukey’s multiple comparison test, ns = P > 0.05, ****P < 0.0001).

### The AGXX^®^ and aminoglycoside synergy exacerbates DNA damage

While our qRT-PCR analysis revealed the transcriptional upregulation of *sulA* in response to AGXX^®^394C/Tob co-treatment (**Fig. 2D**), we next sought to further determine if co-treatment induces significant DNA damage. To monitor and quantify DNA damage, we used *E. coli* cells that overexpress the bacteriophage protein Gam fused to GFP. Gam-GFP has been previously shown to specifically bind to DNA double-strand breaks in both bacterial and mammalian cells (41). Gam-GFP expression was induced by addition of doxycycline in cells either left untreated or treated with AGXX^®^394C and Tob individually or in combination. Only ∼15% and ∼18% of cells treated with Tob and AGXX^®^394C showed one detectable Gam-GFP foci respectively, which did not differ significantly from the ∼5% observed in untreated cells (**Fig. 4A-B**). Combinational AGXX^®^394C/Tob treatment induced a significant increase in Gam-GFP foci, with ∼50% of cells having at least one foci, which was comparable to the positive control treatment with ciprofloxacin, a DNA-damaging fluoroquinolone (**Fig. 4A-B**).

**Fig. 4.**
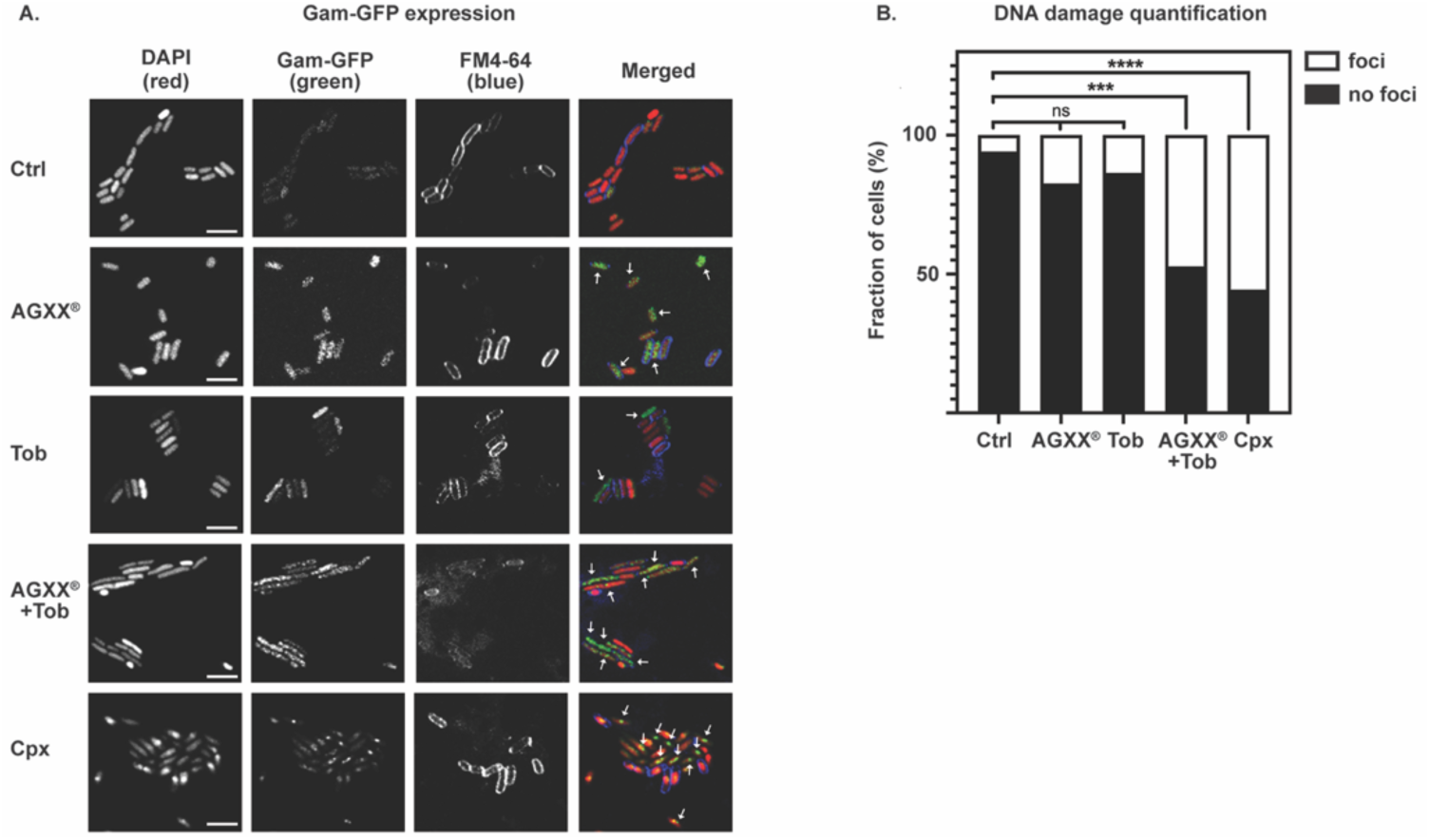
The AGXX^®^ and aminoglycoside synergy exacerbates DNA damage. (**A, B**) Exponentially growing *E. coli* cells expressing Gam-GFP were treated with sub-lethal concentrations of AGXX^®^394C (15 µg/mL), Tob (0.3 µg/mL), or a combination thereof for 3 hours. Untreated cells and treatment with 20 μg/mL ciprofloxacin (Cpx) served as controls. Samples were stained with DAPI (red) and FM4-64 (blue). Confocal images were analyzed by blind-counting Gam-GFP (green) foci. (**A**) Arrows indicate Gam-GFP foci representing sites of DNA damage. Images representative of three biological replicates (scale bar: 5 µm). (**B**) Quantification of the number of foci per cell from microscopy images as shown in **(A)** (n=3; statistics: Two-way ANOVA, Tukey’s multiple comparison test, ns = P > 0.05, ***P < 0.001, ****P < 0.0001).

### The ROS-mediated effects generated by AGXX^®^ and Tob synergy can be suppressed by antioxidants

Our data suggest that the increased sensitivity of UPEC to the synergy between AGXX^®^ and aminoglycosides results from augmented intracellular ROS accumulation, which promotes protein aggregation and DNA damage. It has previously been proposed that the bactericidal effects of aminoglycoside antibiotics are at least in part due to the induction of protein misfolding and insertion of misfolded proteins into the plasma membrane to create membrane channels and uncontrolled drug uptake (6, 42). To this end, we examined whether the known ROS scavenger, thiourea, mitigates the cellular stress induced by AGXX^®^/aminoglycoside combined treatment. To test this, CFT073 cultures were pretreated with thiourea (or left untreated) before exposure to AGXX^®^394C/Tob co-treatment, and bacterial survival and intracellular ROS levels were determined. As expected, in the absence of thiourea, cells subjected to the combined treatment exhibited a ∼3-log reduction in survival (**Fig. 5A**) and a ∼25-fold increase in ROS production (**Fig. 5B**). Thiourea pretreatment restored bacterial survival almost completely, and significantly reduced ROS levels by ∼50% (**Fig. 5A-B**). Furthermore, pretreatment with thiourea also mitigated the induction of IbpA-msfGFP fluorescence to levels observed under unstressed conditions (**Fig. 5C-D**). These findings further suggest that ROS production is a key contributor to the bactericidal effects of the AGXX^®^/aminoglycoside combinational treatment.

**Fig. 5.**
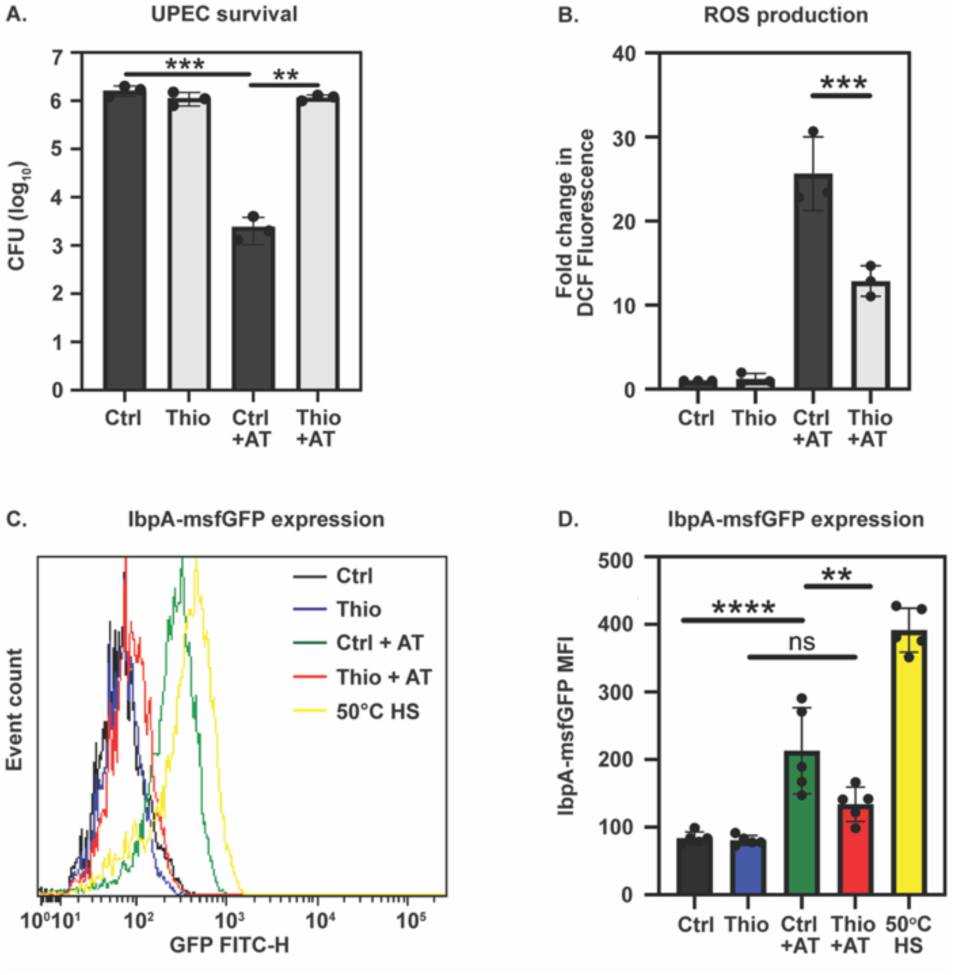
The ROS-mediated synergistic effects between AGXX^®^ and Tob can be suppressed by antioxidants. (**A, B**) Exponentially growing CFT073 cells were exposed for 60 minutes to a combination of AGXX^®^394C (30 µg/mL) and Tob (0.5 µg/mL), labelled as “AT”. Cells left untreated served as the control. Prior to reaching mid-log phase, cells were either left untreated or pretreated with 70 mM thiourea before subsequent exposure to the combined treatment. (**A**) Bacterial survival was assessed by determining CFUs after 1 hour of treatment. (n = 3, ±SD; statistics: One-way ANOVA, Šídák’s multiple comparison test, ** P < 0.01, *** P < 0.001). (**B**) Intracellular ROS levels were quantified using H2DCFDA fluorescence. (n = 3, ±SD; statistics: One-way ANOVA, Šídák’s multiple comparison test, *** P < 0.001). (**C, D**) For IbpA-msfGFP expression analysis via flow cytometry, mid-log phase cells were exposed for 60 minutes to either a combination of AGXX^®^394C (15 µg/mL) and Tob (0.3 µg/mL) (AT), left untreated, or subjected to a heat-shock at 50°C as a positive control. (**C**) Total cellular IbpA-msfGFP fluorescence determined by flow cytometry. One representative dataset of four biological replicates is shown. (**C**) Quantification of mean fluorescence intensity (MFI) of IbpA-msfGFP fluorescence determined by flow cytometry (n = 4, ±SD; statistics: One-way ANOVA, Šídák’s multiple comparison test, ns = P > 0.05, ** P < 0.01, **** P < 0.0001).

### Polyphosphate protects UPEC from AGXX^®^/aminoglycoside co-treatment induced proteotoxicity

In addition to activating protein-based molecular chaperones like IbpA, many Gram-negative pathogens can also respond to proteotoxic antimicrobials by converting cellular ATP into polyphosphate (polyP), a chemical chaperone that aids in preventing protein aggregation (43, 44). We recently showed that UPEC cells lacking the enzyme for polyP production (i.e., Δ*ppk*) are more susceptible to AGXX^®^ (27). To test whether polyP protects UPEC from the proteotoxic effects of AGXX^®^/Tob co-treatment, we analyzed the ability of CFT073 with and without the polyP kinase Ppk1 encoding gene to grow and survive upon exposure to the AGXX^®^394C/Tob combination. A growth comparison of AGXX^®^394C/Tob-treated WT and Δ*ppk* cells showed a significantly longer lag phase of polyP-deficient cells (**Fig. 6A**). Notably, when keeping the concentration of one stressor constant while varying the concentration of the other, it became apparent that polyP protects cells from both AGXX^®^394C and Tob (**Supplementary Fig. S3**). In our survival studies, we exposed exponentially growing WT and Δ*ppk* cells to AGXX^®^394C/Tob for 1 hour, then serially diluted them for CFU counts. In the absence of Ppk1, AGXX^®^394C/Tob co-treatment resulted in a ∼2.5-log lower survival compared to the wild-type, indicating an increased sensitivity to AGXX^®^394C/Tob, which could be fully restored by complementation (**Fig. 6B**). Consistent with these findings, we found that Δ*ppk* cells were also characterized by much higher ROS levels upon treatment with Tob alone and when combined with AGXX^®^394C (**Fig. 6C**). Next, we sought to analyze potential polyP-dependent differences in macromolecular damage using IbpA-msfGFP-expressing WT and Δ*ppk* cells. *E. coli* produces polyP in response to AGXX^®^394C but not to Tob (**Supplementary Fig. 4A-B**). IbpA-msfGFP-expressing WT and Δ*ppk* cells were exposed to a combination of 10 µg/mL AGXX^®^394C and 0.2 µg/mL Tob for 1 hour, which is a combination of AGXX^®^394C and Tob concentrations that resulted in the highest IbpA-msfGFP expression in Δ*ppk* (**Supplementary Fig. 4B-D**). Confocal microscopy showed that IbpA-msfGFP foci in untreated WT and Δ*ppk* cells were comparable with 12% and 6% of all cells, respectively (**Fig. 6D-E**). While in WT cells, the combined treatment caused IbpA-msfGFP foci formation in ∼20% of the cells; foci were counted in ∼85% of Δ*ppk* cells (**Fig. 6D-E**). Activation of the SOS response in bacteria under DNA-damage stress has been shown to block cell division, leading to abnormal growth and filamentation (38, 45). We quantified cell filamentation in AGXX^®^394C /Tob-treated WT and Δ*ppk* cells, which was ∼3-times higher in cells lacking polyP, indicating that DNA damage in polyP-deficient cells is more pronounced (**Fig. 6F**). Flow cytometry analysis further confirmed that AGXX^®^394C/Tob-induced IbpA-msfGFP expression was much more prominent in polyP-deficient cells (**Fig. 6G-H**). Taken together, our data point towards a critical role for polyP in protecting UPEC from the deleterious effects of AGXX^®^/aminoglycoside-induced macromolecular damage.

**Fig. 6.**
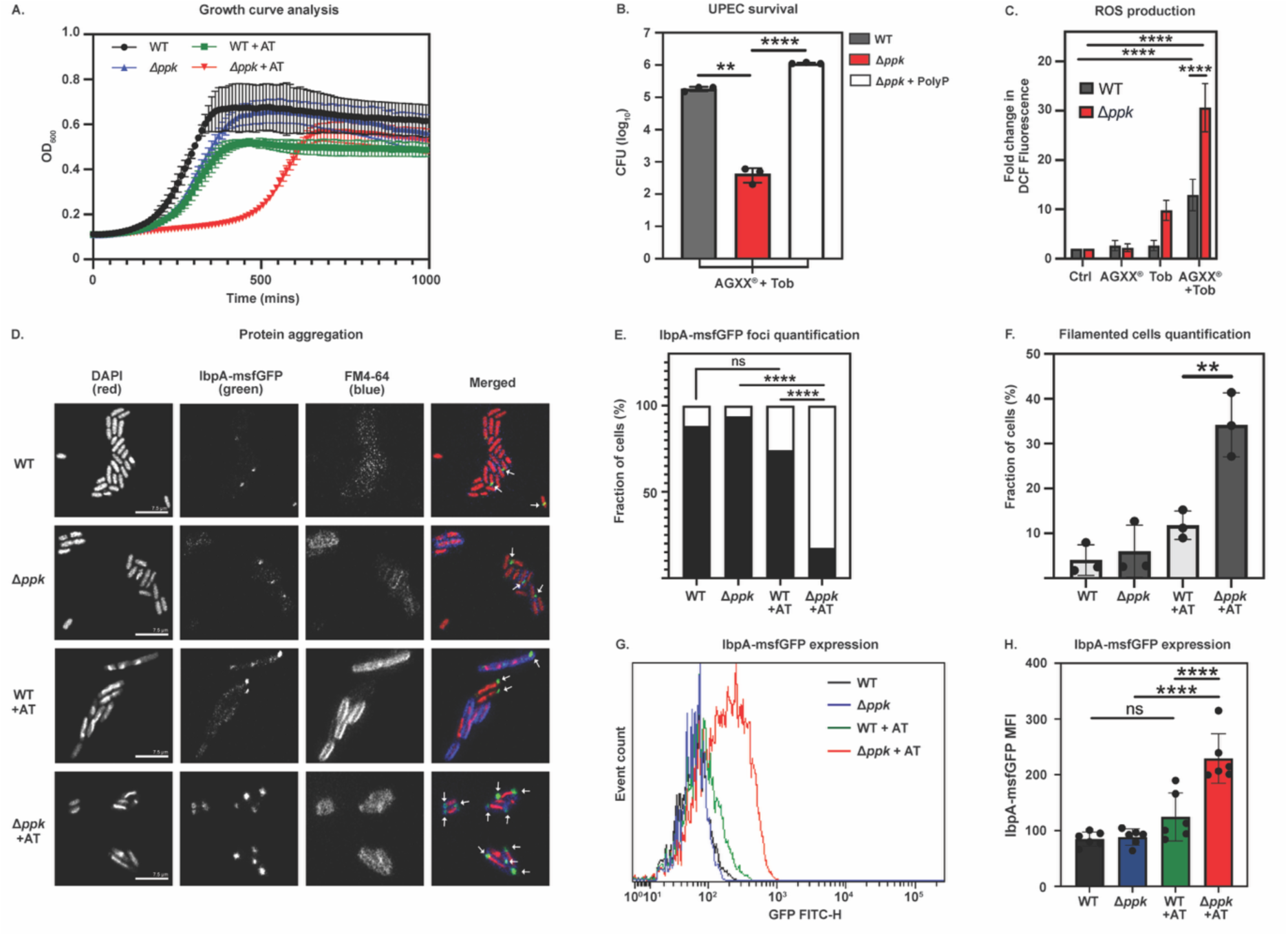
PolyP protects UPEC from AGXX^®^/aminoglycoside-mediated proteotoxicity. (**A**) UPEC strains CFT073 and CFT073Δ*ppk* were treated with a combination of AGXX^®^394C (0.4 µg/mL) and Tob (0.2 µg/mL). Growth was monitored over 18 hours. (n = 3, ±SD). (**B, C**) The indicated strains were treated with a combination of AGXX^®^394C (30 µg/mL) and Tob (0.5 µg/mL) for 60 minutes. (**B**) Survival rate as determined by CFU counts (n = 3, ±SD; statistics: One-way ANOVA, Šídák’s multiple comparison test, ** P < 0.01, ****P < 0.0001). (**C**) Intracellular ROS levels were quantified using H2DCFDA fluorescence (n = 4, ±SD; statistics: One-way ANOVA, Šídák’s multiple comparison test, **** P < 0.0001). (**D-F**) Exponentially growing WT and Δ*ppk* cells expressing IbpA-msfGFP were left untreated or treated with a combination of AGXX^®^394C (10 µg/mL) and Tob (0.2 µg/mL) for 1 hour. (**D**) To visualize and quantify protein aggregates *in vivo*, cells were stained with DAPI (red) and FM4-64 (blue). White arrows indicate foci formed by IbpA-msfGFP (green) binding to protein aggregates *in vivo*. Images representative of three biological replicates (scale bar: 7.5 µm). **(E)** Quantification of the number of foci per cell from three biological replicates (n = 3, ±SD; statistics: Two-way ANOVA, Tukey’s multiple comparison test, ns = P > 0.05, ****P < 0.0001). **(F**) Quantification of the number of filamentous cells observed in microscopy images as represented in **(D)** (n = 3, ±SD; statistics: One-way ANOVA, Šídák’s multiple comparison test, ***P < 0.001). (**G, H**) Total cellular IbpA-msfGFP fluorescence was determined by flow cytometry. (**G**) A representative image from six biological replicates is shown. (**H**) Quantification of mean fluorescence intensity (MFI) of IbpA-msfGFP fluorescence determined by flow cytometry (n = 6, ±SD; statistics: One-way ANOVA, Šídák’s multiple comparison test, ns = P > 0.05, **** P < 0.0001).

## DISCUSSION

### AGXX^®^/aminoglycoside combinations exhibit broad-spectrum antimicrobial activity

The emergence of antibiotic resistance is an escalating global public health concern. Given the lack of new antimicrobials in the industrial pipeline, novel approaches are desperately needed to combat bacterial pathogens that are otherwise difficult to eradicate. The threat of growing antimicrobial resistance is further compounded by the ability of many pathogenic bacteria to form protected multicellular biofilm communities. Not only does biofilm formation provide defense against antimicrobials, which struggle to diffuse through the extracellular matrix shield, but it also enhances the opportunity for the passage of antimicrobial resistance genes between bacterial species via horizontal gene transfer. Biofilm-mediated resistance, along with antimicrobial resistance, is especially problematic in relation to device-associated nosocomial infections (i.e., catheter-associated UTIs, intravenous catheter infection, medical implant-associated infections). To combat these ever-evolving threats, a promising area of research has focused on compounds that enhance the efficacy of antibiotics by targeting metabolic processes and/or cellular networks (46). Compounds with synergistic effects have been reported for aminoglycosides (1), as well as beta-lactams (47–49), rifampicin (50), and macrolides (51). Here, we report on the synergistic interaction between aminoglycosides and AGXX^®^, a ROS-generating surface coating comprised of the two transition metals Ag and Ru. The antimicrobial activity of different AGXX^®^ formulations is attributed to their specific coating composition, including the silver-ruthenium ratio, particle size, and production procedure (27). Silver and silver-containing derivatives such as silver nanoparticles have been previously reported as potent enhancers of aminoglycoside activity in *P. aeruginosa* and other pathogenic bacteria (19, 52–55). These synergies have been shown to have especially meaningful potential clinical implications, with their use as antimicrobial surface coatings on catheters and other medical implants (56). Moreover, silver derivatives such as silver sulfadiazine are currently used in topical applications as a standard for treating and preventing wound infections, despite being associated with side effects such as allergic reactions to the sulfadiazine moiety (57). Intriguingly, AGXX^®^, which has been shown to be significantly more potent than classical silver and silver sulfadiazine (58), is predicted to cause minimal toxicity to human cells due to minimal release of Ag even after several weeks (25, 28), and, to our knowledge, antimicrobial resistance to AGXX^®^ has not been reported yet. Combined, these characteristics make AGXX^®^ potentially well-suited for applications as a novel antimicrobial surface coating for medical devices.

Following up on our previous study, showing that AGXX^®^ potentiates the bactericidal activities of aminoglycosides in the Gram-negative opportunistic pathogen *P. aeruginosa* (26), we now provide evidence that the activity of AGXX^®^/aminoglycoside co-treatments extends to other ESKAPEE pathogens, including MDR-resistant *K. pneumoniae*, *A. baumannii*, *S. aureus*, and UPEC strains (**Fig. 1**; **Fig. 2A**). Our killing assays revealed that, regardless of the strain tested, exposure to combinatorial AGXX^®^/aminoglycoside treatment rapidly causes a 3- to 5-log reduction in bacterial survival compared to the individual treatments. Most importantly, AGXX^®^/aminoglycoside co-treatments reduced the antibiotic concentrations required for efficient killing of clinical ESKAPEE pathogen isolates, which are characterized by high gentamicin and tobramycin resistance, by 8-fold to 16-fold, respectively (**Fig. 1; Supplementary Fig. S1**). This may minimize the risk and/or severity of antibiotic-induced side effects and may also delay or prevent resistance development (59). These data are consistent with our previous observation that AGXX^®^ re-sensitizes an intrinsically resistant *P. aeruginosa* strain to sub-MIC concentrations of kanamycin (26), and underscore the broad-spectrum antimicrobial potential of AGXX^®^.

### The synergy between AGXX^®^ and aminoglycosides is based on the induction of severe oxidative stress that impairs cellular redox homeostasis

Our current study further demonstrates that ROS accumulation is the central mechanism underlying the synergistic interaction between AGXX^®^ and aminoglycoside antibiotics. Consistent with our previous observation in *P. aeruginosa* (26), fluorescence-based assays in UPEC revealed that AGXX^®^/aminoglycoside co-treatments induce a ∼13-fold increase in intracellular ROS levels compared to individual treatments alone (**Fig. 2**). These elevated ROS levels were tightly linked to the induction of canonical stress response pathways, such as the transcriptional upregulation of *ibpA*, which encodes a small heat-shock protein involved in proteostasis, and *sulA*, a marker of SOS response to DNA damage. Thus, our data point towards significant proteotoxic and genotoxic challenges that AGXX^®^/aminoglycoside combinations impose on UPEC, likely exceeding the threshold that a functional redox homeostasis can buffer. Flow cytometry and confocal microscopy further demonstrated that AGXX^®^/Tob co-treatment induced IbpA protein foci to levels comparable to those observed in bacteria under heat shock (Fig. 3). The combinatorial treatment also produced substantial protein aggregation, detected in ∼80% of cells. These observations are consistent with established models in which aminoglycosides promote mistranslation and the accumulation of misfolded proteins (10), while AGXX^®^ independently induces oxidative protein damage through thiol oxidation, leading to widespread protein aggregation (22, 27, 60). As a result, expression of molecular chaperones and proteases is highly upregulated in AGXX^®^-stressed bacteria, as previously shown in several independent transcriptome studies (21–24, 27).

AGXX^®^ has recently been shown to induce detectable DNA double-strand breaks in a ROS-dependent manner, when applied in sufficiently high yet still sub-lethal concentrations (27). In our present study, AGXX^®^/Tob co-treated cells exhibited markedly increased DNA damage, as indicated by the formation of Gam-GFP foci (**Fig. 4**). Gam is a bacteriophage protein that binds to sites of double-strand DNA breaks (41). Although aminoglycosides specifically target protein translation, kanamycin has been reported to damage DNA bases, likely via indirect ROS-mediated damage (61). We did not observe significant Gam-GFP foci formation when cells were treated with AGXX^®^ or tobramycin alone, likely due to the low concentrations of each stressor. Strikingly, co-treatment with AGXX^®^/Tob significantly amplified genotoxic effects, with ∼50% of treated cells displaying Gam-GFP foci and ∼85% of those cells exhibiting multiple foci. This suggests that the combinatorial treatment with AGXX^®^/Tob significantly enhances the DNA-damaging ability of AGXX^®^ in UPEC.

We further demonstrate that ROS are not merely byproducts of, but drivers of, AGXX^®^/aminoglycoside toxicity. We found that pretreatment with the antioxidant thiourea abolished AGXX^®^/Tob-induced ROS accumulation, restored bacterial survival to levels comparable to untreated bacteria, and suppressed IbpA protein foci formation and protein aggregation (Fig. 5). These data therefore establish oxidative imbalance as the initiating event linking AGXX^®^ exposure to enhanced aminoglycoside efficacy. This aligns with ROS-dependent models of aminoglycoside killing (5, 26, 62) and reconciles previous controversies (7, 52) by demonstrating that ROS becomes critical when a redox-active compound amplifies aminoglycoside stress. Together, these ROS-driven insults overwhelm bacterial macromolecules, providing a mechanistic explanation for the pronounced bactericidal synergy of AGXX^®^ with aminoglycosides.

### Bacteria produce polyphosphate to counter AGXX^®^/aminoglycoside toxicity

Providing further evidence to support this ROS-driven bactericidal synergy of AGXX^®^ with aminoglycosides, we found that UPEC protect themselves from this toxicity through polyphosphate (polyP) (**Fig. 6, Supplementary Fig. S4**). Gram-negative bacteria have long been known to produce polyP as a stress defense strategy to counter host defense mechanisms (32, 33, 63–65). In addition to being involved in various biological processes, including biofilm formation (66, 67), motility (68), virulence factor production (69, 70), and host colonization (66, 69, 71), polyP has also been identified as a potent chemical chaperone that protects bacteria from proteotoxic stress (43, 44, 72, 73). Here, we demonstrate a significant increase in polyP production when cells are exposed to AGXX^®^ **(Fig. S4)**. This is consistent with a previous report that showed that *Δppk* cells were highly sensitive to silver nitrate and that the transcriptome of AGXX^®^-treated UPEC cells exhibited signs of silver exposure (27). While the deletion of *ppk1*, the gene responsible for polyP production, has been associated with increased susceptibility of *E. coli* and *P. aeruginosa* towards aminoglycoside and carbapenem antibiotics (74, 75), we did not detect a significant induction in polyP production in Tob-treated cells (**Fig. S4**). Whether the activation of polyP production is in response to AGXX^®^/aminoglycosides-generated ROS, or potentially to silver ions that are released from the AGXX^®^ formulation remains unclear. We conclude that PolyP, likely induced by ROS accumulation, acts as a chemical chaperone to protect from the proteotoxic effects of AGXX^®^/Tob, and propose that both IbpA expression and polyP production are important for a stringent defense. This may explain why, compared to *Δppk* strains, IbpA-msfGFP expression is substantially reduced and IbpA-msfGFP foci are present in much fewer cells in the polyP-producing WT strain. In the absence of polyP, cells depend primarily on IbpA-mediated chaperone activity to counter AGXX^®^/Tob-mediated proteotoxic stress. However, this compensatory mechanism is insufficient to prevent widespread damage, and our data highlights the critical role of polyP in protecting bacterial cells from redox-induced proteome disruptions.

Collectively, our study positions AGXX^®^ as a dual-function antimicrobial that potentiates aminoglycoside effects on Gram-negative pathogens by intensifying ROS-mediated proteotoxic and genotoxic stress. By driving bacterial stress responses beyond recoverable thresholds, AGXX^®^ potentiates the efficacy of aminoglycoside exposure with detrimental effects on bacterial viability. More broadly, our work highlights controlled redox stress as a rational strategy for antibiotic potentiation and provides a mechanistic framework for the development of ROS-based antimicrobial adjuvants.

## MATERIALS AND METHODS

### Bacterial strains and growth conditions

All strains, plasmids, and oligonucleotides used in this study are listed in **Supplementary Table 2.** Unless stated otherwise, overnight cultures were grown aerobically in Luria-Bertani (LB) broth (Millipore Sigma) at 37°C with shaking at 300 rpm for 16-20 hours. For subsequent experiments, overnight cultures were diluted into 3-(N-morpholino) propanesulfonic acid minimal medium supplemented with 0.2% glucose, 1.32 mM K₂HPO₄, and 10 µM thiamine (76), or in artificial urine media (AUM) (29), and incubated at 37°C under shaking conditions.

### Preparation of artificial urine media (AUM)

AUM was prepared as previously described (29). The final concentrations of the components were as follows: 78.5 mM sodium chloride, 9 mM sodium sulfate, 2.2 mM sodium citrate dihydrate, 0.1 mM sodium oxalate, 3.6 mM potassium dihydrogen phosphate, 21.5 mM potassium chloride, 3 g/L tryptic soy broth, 3 mM calcium chloride, 2 mM magnesium chloride, 15 mM ammonium chloride, 6 mM creatinine, and 200 mM urea (pH 6.3). The AUM was freshly prepared every 7 to 10 days.

### Preparation of AGXX^®^ formulations

All AGXX^®^ formulations used in this study were developed and supplied by Largentec GmbH (Berlin, Germany). AGXX^®^394C and AGXX^®^720C microparticles used in this study were made from silver powders on cellulose with a particle size between 1.5 and 2.5 µm (MaTeck, Germany). The silver-ruthenium complex was realized by Largentec proprietary electrochemical deposition processes.

### Time-killing assay of clinical isolates

Overnight cultures were diluted ∼20-fold into MHB by normalizing cultures to an optical density at 600 nm (OD₆₀₀) = 0.1 in a six-well sterile cell culture plate. Samples were then treated with sublethal concentrations of AGXX^®^720C and Gm or Tob individually or in combination. Cell culture plates were incubated at 37°C for four hours at 150 rpm under aerobic conditions. Cells were collected at the indicated time points, serially diluted in phosphate-buffered saline (PBS, pH 7.4), plated on LB agar, and incubated at 37°C for 20 hours to quantify surviving colonies.

### Bacterial survival assays

Overnight cultures of UPEC strains CFT073 or CFT073Δ*ppk* were diluted 50-fold into the indicated media to an initial OD₆₀₀ ∼ 0.08 and incubated until reaching an OD₆₀₀ = 0.3. Cultures were then transferred to sterile 125 mL flasks and exposed to sublethal concentrations of AGXX^®^, the indicated aminoglycosides, or both. If indicated, cultures were pretreated with 70 mM thiourea for 60 minutes before AGXX^®^ and/or aminoglycoside exposure. After 1 hour, aliquots were serially diluted in PBS, plated on LB agar, and incubated overnight at 37°C with shaking to determine colony-forming unit (CFU) counts.

### Checkerboard Assay

Checkerboard assays were performed as previously described (77), with slight modifications. 100x stock solutions of the indicated aminoglycoside antibiotics were serially diluted and added to the respective wells of 12-well plates, while the concentration of AGXX^®^720C was held constant across wells. Exponentially growing UPEC CFT073 cultures in MOPSg minimal media were adjusted to an initial OD₆₀₀ of approximately 0.05 and added to each well. Plates were incubated at 37°C with shaking for 16–18 hours. OD₆₀₀ was subsequently measured to assess bacterial growth.

### Intracellular ROS quantification

Intracellular ROS levels were quantified using the redox-sensitive dye 2′,7′-dichlorodihydrofluorescein diacetate (H₂DCFDA) (Thermo Fisher Scientific). Exponentially growing cultures of CFT073 and CFT073Δ*ppk* were left untreated or treated with the indicated concentrations of AGXX^®^394C, Tob, or their combination for 1 hour. Duplicate samples were then collected, washed once with pre-warmed PBS, normalized to an OD₆₀₀ ∼1.0 in prewarmed PBS containing 10 µM H2DCFDA, incubated in the dark at 37°C for 30 minutes, rewashed with PBS before DCF fluorescence was measured at excitation/emission wavelengths of 485/535 nm in a Tecan Infinite 200 plate reader. If indicated, cells were incubated with 70 mM thiourea, a ROS quencher, prior to stress treatments.

### Gene expression analysis by qRT-PCR

Overnight cultures of CFT073 were diluted into MOPSg medium to an OD₆₀₀ ∼ 0.08 and grown to mid-log phase (OD₆₀₀ ∼ 0.3). Cultures were either left untreated or treated with 30 μg/mL AGXX^®^394C, 0.5 μg/mL Tob, or their combination. Following 10 mins of incubation, transcription was halted by the addition of ice-cold methanol. Total RNA was extracted from at least three biological replicates using a commercial RNA extraction kit (Macherey-Nagel), residual genomic DNA was removed with the TURBO DNA-free kit (Thermo Scientific), and mRNA was reverse transcribed into cDNA using the PrimeScript cDNA synthesis kit (Takara). qRT-PCRs were performed using manufacturer protocols (Alkali Scientific) with gene-specific primers (e.g., *ibpA, sulA*, and *rrsD*). Gene expression was normalized to *rrsD*, and relative transcript levels were determined using the 2^−ΔΔCT^ method (78).

### IbpA-msfGFP expression by flow cytometry

Overnight cultures of strains MG1655::*ibpA-msfGFP* and MG1655Δ*ppk::ibpA-msfGFP* were diluted into fresh MOPSg medium to an OD₆₀₀ ∼0.01 and grown to mid-log phase (OD₆₀₀ ∼ 0.3). Prior to cultures reaching mid-log phase, cells were then left untreated or pretreated with 70 mM thiourea before the indicated concentrations of AGXX^®^394C, Tob, or their combination were added for 1 hour. Cells exposed to 50°C for 60 mins served as positive controls. For flow cytometry analysis of IbpA-sfGFP expression, samples were normalized to an OD₆₀₀ of 0.05 in PBS (pH 7.4) and analyzed on a BD FACS Melody cytometer via the FITC channel, recording at least 10,000 events per sample. Data were analyzed with FCSalyzer.

### IbpA-msfGFP binding to protein aggregates *in vivo*

Overnight cultures of strains MG1655::*ibpA-msfGFP* and MG1655Δ*ppk::ibpA-msfGFP* were diluted into fresh MOPSg medium to an OD₆₀₀ ∼ 0.01 and grown to mid-log phase (OD₆₀₀ ∼ 0.3). Prior to cultures reaching mid-log phase, cells were then left untreated or pretreated with 70 mM thiourea before the indicated concentrations of AGXX^®^394C, Tob, or their combination were added for 1 hour. Cells exposed to 50°C for 60 min served as positive controls. For visualization of IbpA-sfGFP-bound protein aggregates, cells were washed twice in PBS, stained with 5 µg/mL DAPI and 5 µg/mL FM4-64 in the dark for 15 minutes, rewashed, and mounted on 1% agarose pads. Fluorescence microscopy was performed using a Leica SP8 confocal system with a DMi8 CS inverted microscope. At least 500 cells across all replicates were blindly quantified.

### Gam-GFP binding to DNA double-strand breaks

Strain MG1655::*gam-gfp* was grown overnight in LB supplemented with 200 ng/mL doxycycline to induce Gam-GFP expression. Cells were diluted into doxycycline-containing MOPSg medium to an OD₆₀₀ ∼0.08 and grown to mid-log phase (OD₆₀₀ ∼ 0.3), and treated with the indicated concentrations of AGXX^®^394C, Tob, or their combination. The addition of ciprofloxacin was used as a positive control for DNA damage. Following 3 hours of treatment, cells were washed twice, resuspended in PBS, stained with 5 µg/mL DAPI and 5 µg/mL FM4-64 in the dark for 15 minutes, rewashed and resuspended in PBS. Samples were imaged on 1% agarose gel pads using a Leica SP8 confocal system, which is equipped with a DMi8 CS inverted microscope. At least 500 cells across all replicates were counted blindly.

### Construction of in-frame gene deletions in *E. coli* cells expressing IbpA-msfGFP

An in-frame *ppk* deletion mutant was constructed using the lambda red-mediated site-specific recombination (79). In brief, the *ppk* gene in *E. coli* MG1655-*ibpA*-msfGFP was replaced by a chloramphenicol resistance cassette, which was subsequently resolved using pCP20 to yield the nonpolar in-frame deleted strain MG1655-*ibpA*-msfGFPΔ*ppk*. The chromosomal mutation was verified by PCR.

### PolyP extraction and quantification

Intracellular polyP levels were extracted as previously described (70, 80), with minor modifications, and quantified using DAPI. Mid-log-phase cultures of the IbpA-msfGFP-expressing *E. coli* WT and Δ*ppk*1 strains were grown in MOPSg at 37°C under aerobic conditions and then exposed to the indicated concentrations of Tob or AGXX^®^394C for 1 h. Following treatment, cultures were normalized to an OD₆₀₀ of 0.3, and 1mL aliquots were harvested by centrifugation at 8,000 x g, 5 min, 4°C. Cell pellets were resuspended in 200 µL guanidinium isothiocyanate (GITC) lysis buffer and incubated at 95°C for 10 min to release intracellular polyP. A 50 µL aliquot of the crude lysate was reserved for total protein determination using the Bradford assay. The remaining lysate was mixed with 95% ethanol and applied to silica membrane EconoSpin columns (Epoch Life Science) for polyP purification. Columns were washed twice with 5 mM Tris-HCl (pH 7.5), and polyP was eluted with 50 mM Tris-HCl (pH 8.0). Eluted polyP was mixed with 25 μM DAPI, incubated in the dark for 10 min, and fluorescence was measured at excitation/emission wavelengths of 420/550 nm, which preferentially detects DAPI–polyP complexes. DAPI in the elution buffer was deducted as the buffer control. Further, the background was determined by treating the purified polyP samples with 1 mg Ppx for an hour prior to staining with DAPI. PolyP concentrations were then extrapolated using a sodium polyP standard curve that was prepared in 50mM Tris HCl, pH 8.0.

### Growth curve-based assays

Overnight cultures of CFT073 and CFT073Δ*ppk* were diluted ∼50-fold into MOPSg media, grown to early stationary phase, diluted again into fresh MOPSg to an OD_600_ = 0.02, and incubated in a Tecan Infinite 200 plate reader with or without the indicated concentrations of AGXX^®^394C, Tob, or their combination. Absorbance at 600 nm was measured every 10 minutes for 16 hours.

### Statistical analyses

All statistical analyses were performed in GraphPad Prism version 8.0.

## ACKNOWLEDGEMENTS

This work was supported by the NIAID grant 1R03AI174033-01A1 (to J.-U. D.). We thank members of the Dahl and Floyd labs for discussion and feedback. The Largentec GmbH team is acknowledged for providing the different AGXX^®^ formulations and for helpful discussions. Susan Rosenberg (Baylor College of Medicine) and Abram Aertsen (KU Leuven) are acknowledged for strains expressing Gam-GFP and IbpA-msfGFP, respectively.

## ETHICS STATEMENT

This work did not involve human subjects or vertebrate animals.

## AUTHOR CONTRIBUTIONS

Emmanuel P. Oladokun: investigation, visualization, writing first draft; Gracious Y. Donkor: investigation, review and editing; Julius K. Narh: investigation; Grady D. Jacobson: investigation; Cade Ward: investigation; Patrick O. Tawiah: investigation, review and editing; Lisa K. Polzer: investigation; Kevin A. Edwards: methodology, review and editing; Kyle A. Floyd: review and editing; and Jan-Ulrik Dahl: conceptualization, supervision, funding acquisition, writing – review and editing.

## DATA AVAILABILITY STATEMENT

Data available on request from the authors.

