## Supplementary Figures & Tables for "The Synergy between a Silver-Ruthenium Antimicrobial and aminoglycosides is based on severe macromolecular damage"

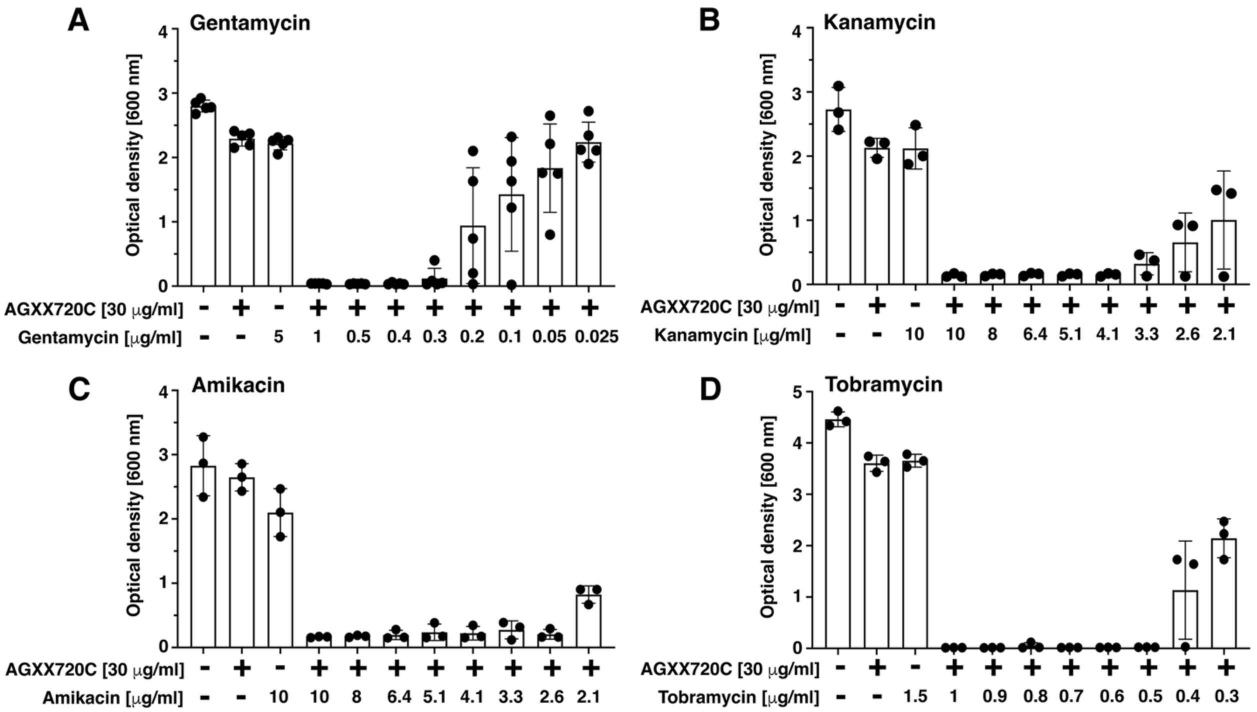


**Supplementary Figure S1.** The impact of the antibacterial activity of gentamycin (**A**), kanamycin (**B**), amikacin (**C**), and tobramycin (**D**) in the presence and absence of 30 μg/mL AGXX®720C was assessed using a checkerboard assay. The $OD_{600}$ was measured after 16-18 hours of incubation. Each plot shows the $OD_{600}$ of *E. coli* CFT073 cells in the presence of varying concentrations of aminoglycosides, with or without AGXX®720C. (n = 3-5, ±SD).

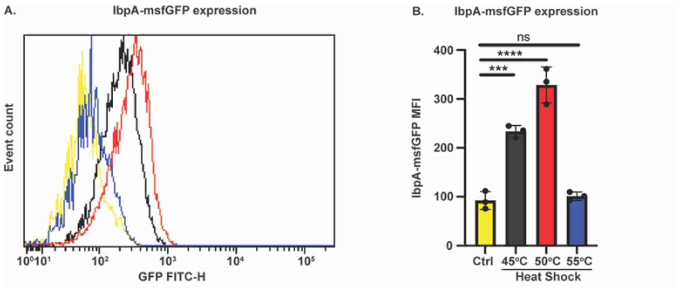


**Supplementary Figure S2:** Cellular IbpA-msfGFP fluorescence was quantified in mid-log phase *E. coli* cells subjected to heat-shock at the indicated temperatures for 60 min. (**A**) Cellular IbpA-msfGFP fluorescence was determined by flow cytometry. One representative dataset from three biological replicates. (**B**) Mean fluorescence intensity (MFI) of IbpA-msfGFP fluorescence from three biological replicates was quantified.

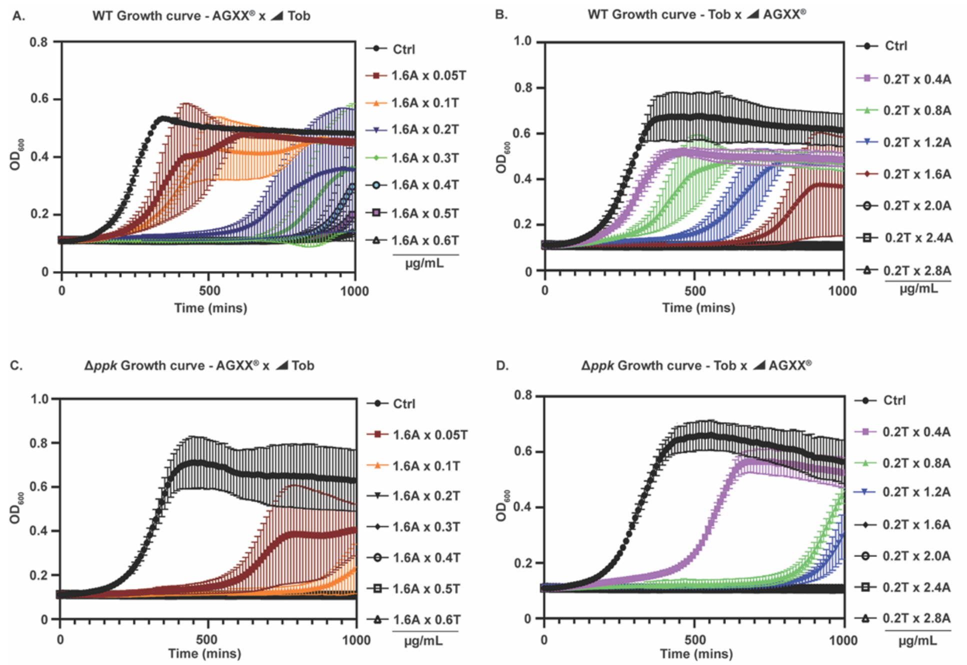


**Supplementary Figure S3:** UPEC strains CFT073 and CFT073Δ*ppk* were diluted into MOPSg to an OD600 = 0.02, treated with or without a combination of AGXX®394C and Tob at the indicated concentrations, and incubated for 18 hours to monitor growth. (**A&C**) WT (**A**) and Δ*ppk* (**C**) treated with 1.6 μg/ml AGXX®394C and varying Tob concentrations. (**B&D**) WT (**B**) and Δ*ppk* (**D**) treated with varying AGXX®394C concentrations and 0.2 μg/ml Tob. (n = 3, ±SD).

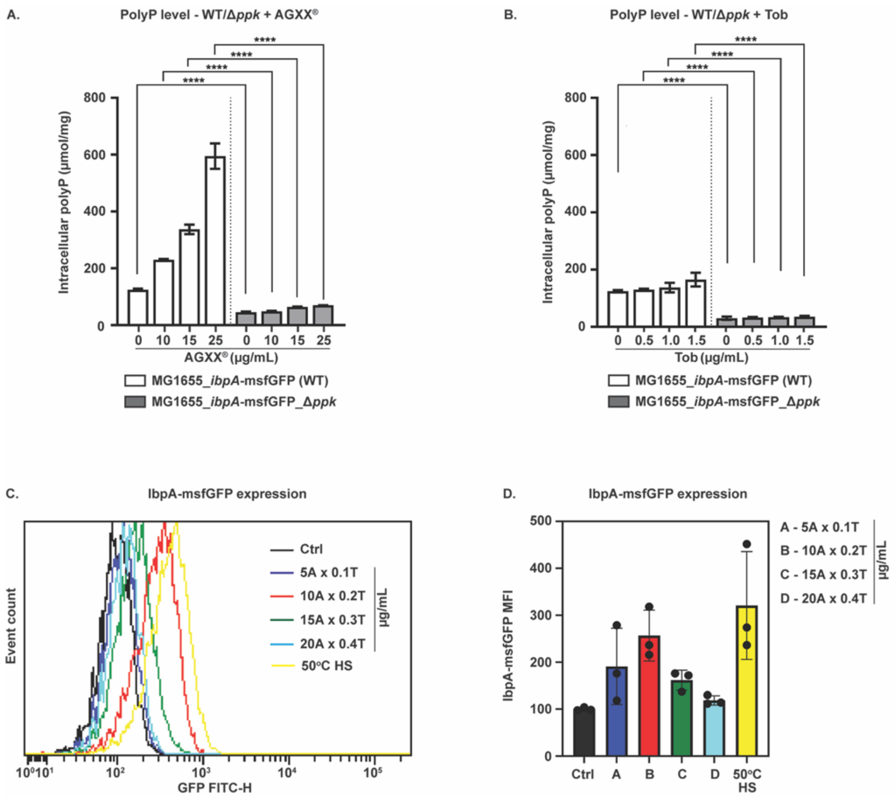


**Supplementary Figure S4:** (**A, B**). Intracellular PolyP was extracted from exponentially growing WT and Δ*ppk* cultures that were treated with and without the indicated concentrations of (**A**) AGXX®394C and (**B**) Tob. Extracted polyP was quantified with 25 µM DAPI. All samples were also treated with 1 mg Ppx and deducted from the RFU as background. DAPI-polyP fluorescence was measured at excitation/emission wavelengths of 415 and 550 nm, respectively. PolyP concentrations were calculated using a sodium polyP standard curve; (n = 3, ±SD). Statistical analysis was performed using one-way ANOVA (n = 3, ±SD; ****P < 0.0001). (**C, D)** Exponentially growing WT cells expressing IbpA-msfGFP were left untreated or treated for 1 hour with the AGXX394C/Tob combination at the indicated concentrations. Cells heat-shocked at 50 ˚C for 60 min served as controls. Cellular IbpA-sfGFP fluorescence was quantified by flow cytometry. (**C**) A representative image from three biological replicates is shown and (**D**) MFI of IbpA-sfGFP was quantified.

**Supplementary Table S1: Minimal inhibitory concentrations (MICs) of the indicated stains**

| Strain/Organism | MIC | |
|---|---|---|
| | **Gentamicin (µg/ml)** | **Tobramycin (µg/ml)** |
| *Staphylococcus aureus* (Strain 3) | 1 | 1 |
| *Klebsiella pneumoniae* (Strain 18) | 1.25 | 1.25 |
| *Acinetobacter baumannii* (Strain 99) | >2000 | 500 |
| *Pseudomonas aeruginosa* (Strain 1) | >2000 | >2000 |
| *Pseudomonas aeruginosa* (Strain 36) | 10 | 2.5 |
| *Pseudomonas aeruginosa* (Strain 40) | 10 | 2.5 |

**Supplementary Table S2: Strains, plasmids, and Oligos used in this study**

| Strain name | Relevant Genotype/Characteristics | Antibiotic Marker |
|---|---|---|
| MG1655-*ibpA-msfGFP* | MG1655 *ibpA-msfgfp* | |
| MG1655Δ*ppk-ibpA-msfGFP* | MG1655Δ*ppk-ibpA-msfGFP* | |
| MG1655-gam-GFP | MG1655 Δ*araBAD567* Δ*attλ::PBAD zfd2509.2::PN25tetR FRT* Δ*attTn7::FRTcatFRT PN25tetOgam-gfp* | |
| CFT073 | UPEC reference strain | |
| VUTI207 | UPEC clinical isolate strain 207 from the Vanderbilt University Medical Center collection | |
| VUTI156 | UPEC clinical isolate strain 156 from the Vanderbilt University Medical Center collection | |

| Plasmid | Description | Antibiotic Marker |
|---|---|---|
| pKD3 | *cat*$^{+}$ chloramphenicol resistance cassette donor | Cm$^{R}$ |
| pKD46 | λ Red recombinase$^{+}$ | Amp$^{R}$ |
| pCP20 | Flp recombinase$^{+}$ | Amp$^{R}$ |

| Primer | Sequence (5'-3') |
|---|---|
| *ppk*-Del-fw | CGC CAT AAT ATC CAG GCA GTG TCC CGT GAA TAA AAC GGA GTA AAA GTG GTA ATG GTG TAG GCT GGA GCT GCT TC |
| *ppk*-Del-rev | ACC GCA GCA AAC TCC TGC GGA CGA GGG GAT TTA TCG TGT ATT GGC ATA GGG TTA CAT ATG AAT ATC CTC CTT AG |
| Δ*ppk*-fw | CCT GTA AAT CGC AAG CTC CA |
| Δ*ppk*-int_fw | AAC CCA TAC CGT CCG GTG AC |
| Δ*ppk*-rev | TTG CCT CTT CAC TCA ACA TA |
| *sulA*_Fw | GGCTTATCAGTGAAGTTGTC |
| *sulA*_rev | TGTTGCGGTGTTAACCAGAG |
| *ibpA*_Fw | ACAACCAGAGCCAGAGTAATGG |
| *ibpA*_rev | TATCCTGGGCGGTAATTTCC |
| *rrsD*_fw | agaaccttacctggtcttgacatc |

| | |
|---|---|
| *rrsD*-rev | cagtttatcactggcagtctcctt |
